# Single-cell transcriptome lineage tracing of human pancreatic development identifies distinct developmental trajectories of alpha and beta cells

**DOI:** 10.1101/2021.01.14.426320

**Authors:** Li Lin, Yufeng Zhang, Weizhou Qian, Yao Liu, Yingkun Zhang, Fanghe Lin, Cenxi Liu, Guangxing Lu, YanLing Song, Jia Song, Chaoyong Yang, Jin Li

## Abstract

In comparison to mouse, the developmental process of human islets has not been properly elucidated. The advancement of single cell RNA-seq technology enables us to study the properties of alpha and beta cells at single cell resolution. By using mitochondrial genome variants as endogenous lineage-tracing markers, we found that human alpha and beta cells have different lineage features. This finding suggests specific endocrine progenitors for alpha and beta cells, which is different from mouse islet cells. This strategy was also applied to a study of chemically-induced islet cell reprogramming and was used to help identify artemether-induced alpha-to-beta trans-differentiation in human islets. The computational results of this study will inspire future studies to establish, maintain, and expand beta cell-specific progenitors *in vitro* and *in vivo*.

## Introduction

Pancreatic islets are composed of a few distinct cell types, including alpha, beta, delta, pp, and ghrelin cells. With mice serving as a model organism, Cre-LoxP based lineage-tracing systems have previously been employed by developmental biologists to characterize pancreatic islet development in mammals [1–5]. One important feature of mice islet development is that the alpha and beta cells share the same endocrine progenitor. Currently, there is a lack of evidence for the existence of cell-type-specific endocrine progenitors [6, 7].

Whether this feature of alpha and beta cells also exists in the development of human islets has not been tested to date. Unlike mice models, it is nearly impossible to study human islet development *in vivo* with lineage-tracing systems. However, the development of single cell RNA-seq (scRNA-seq) technology can shed light on solving the mystery of human islet development [8–10]. The advancement of computational algorithms such as pseudo-time analysis could provide us with tools to understand such developmental processes [11–13]. Unfortunately, these computational tools can only provide a hypothesis based on expression profiles, but not lineage-tracing evidence.

The mutation of the mitochondrial genome in progenitors may be inherited by mature cell types subjected to this development process. Thus, mitochondrial genome variants can be used as an endogenous lineage-tracing marker to study cells [14, 15]. Such an analysis normally requires single cell ATAC-seq or RNA-seq data of cells in different developmental stages. Unfortunately, these data are particularly difficult to obtain from tissues like human islets *in vivo*.

In the current manuscript, we designed a computational method to describe the lineage features and predict the developmental path of mature human islet cells. The mitochondrial mutation sites, which are either group markers between two different cell types or contain high variance among all cell types, are called in order to infer clustering and lineage information for human islet cells (**Fig. 1A**). To deal with different datasets that resulted from various scRNA-seq platforms, including both full-length scRNA-seq and end-counting methods, we developed two procedures to analyze cell-level and cell group-level inferences. Using our method, we identified the potential existence of cell-type-specific progenitors in human islet differentiation processes. In addition, we applied our method to validate the chemically-induced transdifferentiation of alpha cells to beta cells in primary human islets.

**Figure 1.**
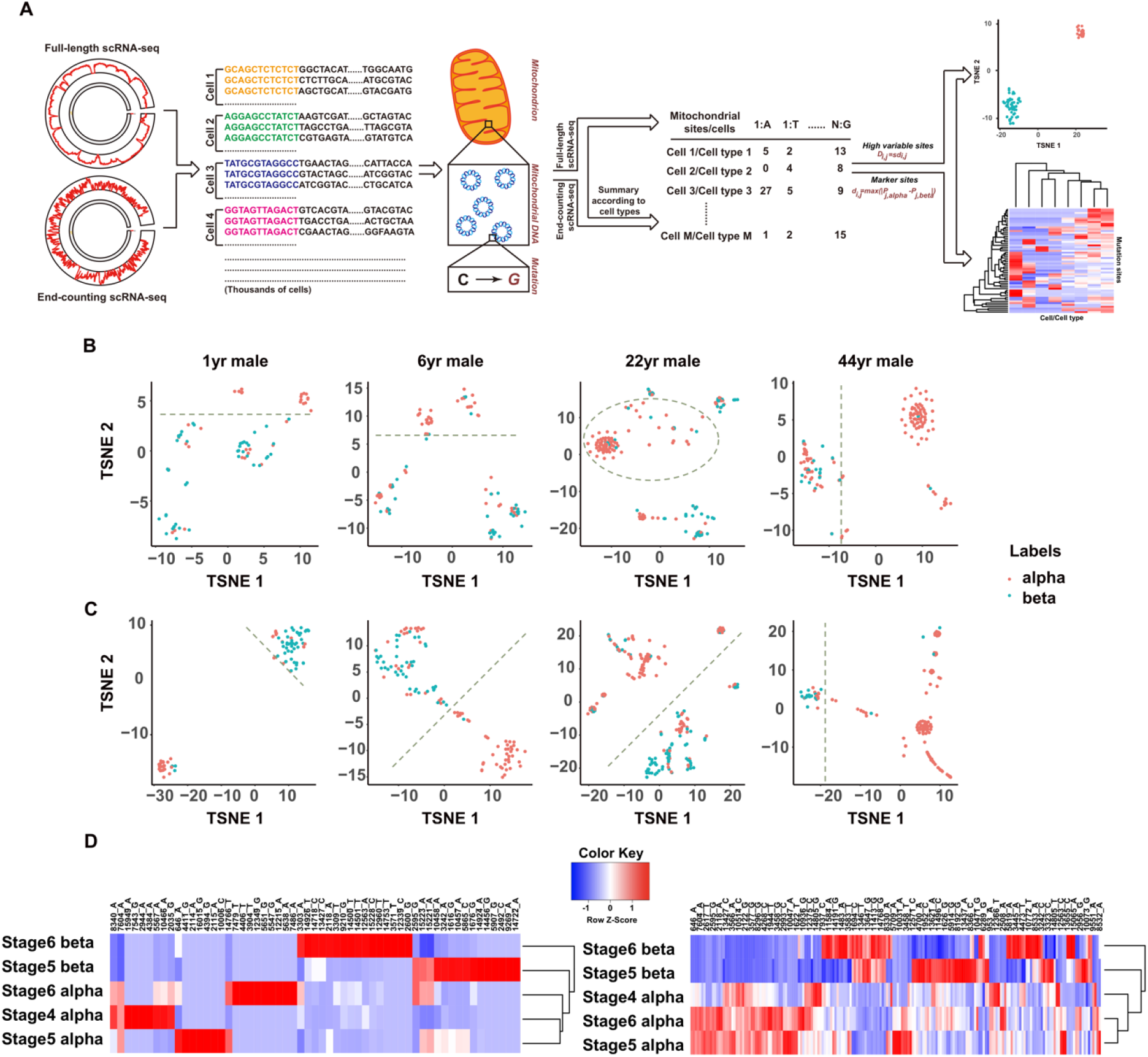
Clustering of alpha and beta cells based on lineage information. **A.** Schematic overview of our lineage-tracing analysis with mitochondrial genome variants based on scRNA-seq data. **B**. Clustering of primary human islet cells based on 10 sites with the highest variants among all cells. **C.** Clustering of primary human islet cells based on top 10 marker sites. **D.** Clustering of iPS cell-derived human islet cells based on high variance sites (left) and marker sites (right) respectively.

## Results

### Human alpha and beta cells show different lineage features

To explore the lineage features and predict the developmental paths of human alpha and beta cells, we first employed Smart-Seq2 (SS2) data derived from human islets from donors at various ages [16]; and secondly, inDrops data of Induced pluripotent stem cell (iPS) cell-derived human islet cells [8]. Due to the low sequence coverage of the inDrops data, cells annotated as the same type were analyzed as one sample. The basic information of these datasets, including the number of cells assigned to different cell types, was summarized and is presented in **Supplementary Table 1 and Table 2**. The single nucleotide variants of the mitochondrial genome were called by comparing the aligned reads with the reference genome. The properties of mutations used in the clustering are summarized in **Supplementary Table 3 and Table 4**. We found that there were both nonsense and missense mutations within the top variants. The mutation burden in alpha and beta cells for both primary and iPS cell-derived cells are shown in **Supplementary Figure 1**.

For the SS2 data, the lineage-based clustering of these cells was executed using two different strategies and visualized based on t-distributed stochastic neighbor embedding (t-SNE). For clustering, we used the top 10 variance mitochondrial genome sites, among cells, firstly, and then specifically between alpha and beta cells (defined as group marker sites). Surprisingly, we were able to differentiate alpha cells from beta cells easily based on their mitochondrial genome mutation information given either of these strategies (**Fig. 1B and C**). More importantly, the clustering results were not affected by the age of donors, but rather the amount of alpha and beta cells represented in the dataset (**Supplementary Figure 2**). We hypothesize that this clustering might not work properly if there are vastly more cells of one cell type than the other. Similar results were identified on the iPS cell-derived human islet cells (**Fig. 1D**). Based on the distinct lineage features between alpha and beta cells, we predict that alpha and beta cells develop from different progenitors.

### Validation of alpha-to-beta transdifferentiation via chemical treatment on human islets

The discovery of cell-type-specific lineage features also allows the lineage-tracing study of mature human islet cells, which have undergone reprogramming. The inducible Cre-LoxP system has been widely used to study islet cell reprogramming in mice [17–19]. However, such a system is difficult to apply to intact primary human islets. For example, a previous study showed that artemether treatment increased the expression of beta cell-specific genes in human alpha cells [20]; but, the so-called artemether-treated “alpha cells” were assigned as alpha cells purely based on their transcriptomes instead of lineage-tracing markers. Therefore, we used mitochondrial genome variants to perform lineage-tracing studies on scRNA-seq data.

The lineage analysis was performed on mitochondrial genomes specifically between alpha and beta cells before artemether treatment from the same donors. The variants are visualized in **Fig. 2A**. K-means clustering was performed based on the six group marker sites, which show a frequency difference greater than 0.05 among all of the five paired alpha and beta drug-untreated samples. We identified a much clearer separation between alpha and beta cells, which might be due to a spike-in of artificial RNA in this dataset to decrease the noise from cell-free RNA (**Fig. 2B**). Based on the six marker sites, alpha cells with or without artemether groups clustered together, indicating that the artemether-treated alpha cells were derived from alpha cells and not beta cells. In addition, the group marker sites are shown in **Fig. 2C**. The lineage features of artemether-treated alpha cells were very similar to the DMSO-treated alpha cells, but not the beta cells.

**Figure 2.**
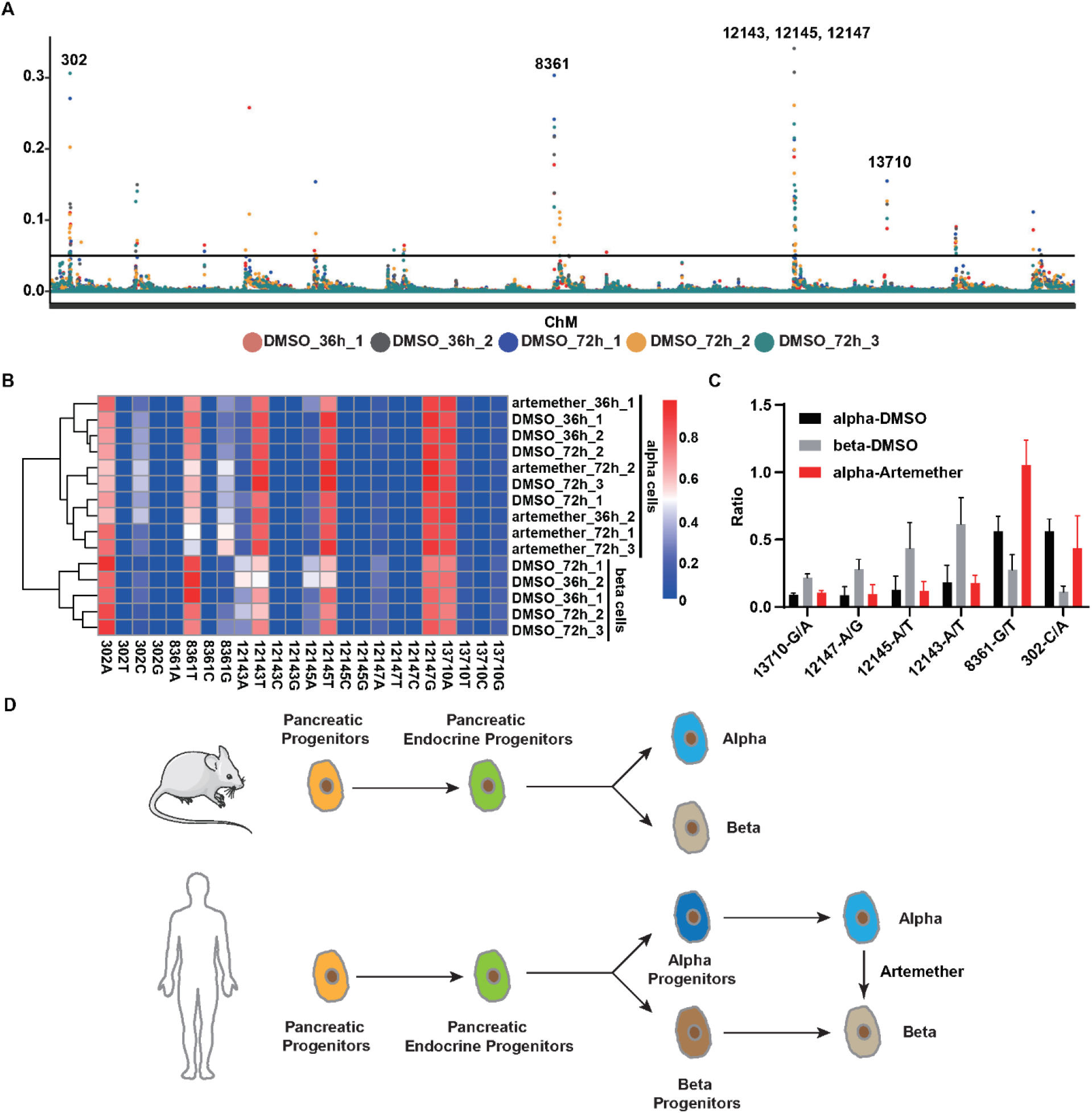
Lineage-tracing study of artemether induced alpha-to-beta transdifferentiation. **A.** Cell-type-specific mitochondrial genome variations were identified by calculating base frequency differences among pairs of alpha and beta cell samples. **B.** Unsupervised hierarchy clustering was performed with genotype variation. Colors indicate the frequency of corresponding genotypes within the site. **C.** The level of major variation was shown as the ratio between two bases in the group marker sites. **D.** Summary of the study. Human alpha and beta cells might develop from different progenitors, and artemether induces alpha-to-beta transdifferentiation in primary human islets.

## Discussion

The differences between mice and human tissues, including cellular composition and functions, have been appreciated for a long time. However, it has never been studied whether human islet cells have a significantly different *in vivo* developmental pathway in comparison to mice. Our study, for the first time, provides computational evidence of cell-type-specific lineage features on primary human islets and iPS cell-derived human islet cells. We found that human alpha and beta cells show distinct mitochondrial variants of scRNA-seq data from two different platforms in multiple donors. These findings also enable us to computationally validate artemether-induced alpha-to-beta transdifferentiation. Therefore, we hypothesize that (1) human alpha and beta cells develop from different progenitors; and (2) artemether induces alpha-to-beta trans-differentiation in primary human islets (**Fig. 2D**).

The existence of alpha and beta cell-specific progenitors has long been speculated. *In vitro*differentiation assay reveals the existence of an Nkx6.1+ stage between endocrine progenitors and mature beta cells [8]. Single cell RNA-seq studies on the fetal pancreas of 9 weeks of development have identified progenitors with potential correlations to either alpha or beta cells [9]. However, these studies do not provide any lineage information. We are aware that there is a lack of biochemical data from wet lab experiments in our current study and that these findings will need verification in the future. However, our study provides clear lineage information based on computational analysis of RNA-seq data and will serve to inspire research in the scientific community. The potential existence of cell-type-specific progenitors makes it meaningful to establish, maintain, and expand beta cell-specific progenitors *in vitro* and *in vivo*. These cells, which might differentiate into functional beta cells upon induction, can serve as an important resource for not only biomedical research but also cell therapies for insulin-deficient patients.

## Methods

### Data processing

All pancreatic islet datasets used in this study were obtained from the following accession numbers: GEO: GSE81547(Smart-Seq2), GEO: GSE114412 (Stages 3 to 6 with Protocol x1 under GSM3141954-3141957, inDrops), GEO: GSE147203 (only human islet data is used, 10x) and were aligned to the GRCh38 human genome and its associated annotations (GRCh38.98). For the Smart-Seq2 [16], the reads were aligned using STAR version 2.7.1a with default parameters. For inDrops[8], raw fastq reads were processed according to the custom in Drops pipelines (www.github.com/indrops/indrops) using Trimmomatic to trim reads, Bowtie 1.2.3 to map reads to the reference transcriptome, and the parameters were those in the default_parameter YAML file. For 10x [20], the default cellranger [15] pipeline was applied.

### Mitochondrial genotype matrix generation

MtDNA reads were extracted with samtools (version 1.9) from the alignment results to generate a new bam file. Then, for each individual bam file, the total number of reads aligned per allele was counted using a Python script[15]. In brief, aligned reads were filtered by the minimum alignment quality of 30 and the minimum mean base-quality of 30. The allele frequency (AF) of a base b at a specified position x on the mitochondrial genome was computed as:

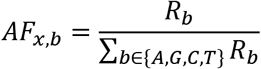

where *R_b_* is the number of reads holding the base b at position x; and ∑_*b*∈{*A,G,C,T*}_ *R_b_* is the coverage of position x.

For Smart-Seq2, a mitochondrial genotype matrix was generated by taking all the A, G, C, T variations for single cells in which a row represents each cell and a column represents each gene. Owing to the low sequence coverage of inDrops and 10x, cells annotated as the same cell type were processed as one sample. The assembled sample data were analyzed with the same pipeline as for Smart-Seq2 data processing in order to extract a mitochondrial genotype matrix with a row corresponding to each cell type and a column for each gene.

### High variance site and group marker site identification

High variance site calling was conducted by identifying the sites with specific genotypes that showed the highest standard deviation (SD) among cells/samples. Specifically, we defined 10 sites with the highest SDs as high variance sites for datasets generated from full-length scRNAseq methods (eg: Smart-Seq2). Sites with a SD greater than 0.3 among all samples were defined as high variance sites for datasets generated from end-counting methods (eg: 10x, inDrops). To screen the marker sites, which show high variance among different cell types, receiver operating characteristic curves (ROC curves) were built based on the comparison between genotype frequency in each site and cell type annotations. Ten sites with the highest AUC (the area under the ROC curve) scores were defined as marker sites for datasets generated from full-length scRNAseq methods. For datasets generated from end-counting methods, the sites that showed phenotype frequency differences greater than 0.05 between all predefined paired samples were defined as marker sites. Finally, alpha and beta samples (merged from cells) from stage 5/6 and the drug-untreated stage of each donor were used as predefined paired samples for the inDrops dataset and 10x datasets, respectively.

### Clustering

For Smart-Seq2 data, we performed a 2D t-SNE visualization based on the high variable sites and marker sites, respectively. For inDrops and 10x data, hierarchical clustering was applied to infer the group and lineage information among cell types based on high variance sites and marker sites, respectively.

### Genetic variant annotation

To further explore high variance and marker sites, SnpEff v5.0 (http://pcingola.github.io/SnpEff/) was used to finish the genetic variant annotation and functional effect prediction.

## Supporting information

Supplemental information of the manuscript

## Data availability statement

The source code has been deposited to github. The link can be provided upon request and will be open to public once the manuscript gets accepted.

## Acknowledgments

This work was supported by China National Key Research and Development Program 2018YFA0801300, 2020YFA0803600, and China National Natural Science Foundation 32071138 to J.L.

## Author contributions

Conceptualization, J.S. and J.L.; Investigation, L.L., Y.Z., W.Q., Y.L., Y.Z., F.L., C.L., G.L., Y.S.; Writing, J.S. and J.L.; Data Visualization, L.L., Y.Z., and W.Q.; Funding Acquisition, J.L.; Supervision, C.Y. and J.L.

## REFERENCE

1. Schindehutte J, Fukumitsu H, Collombat P, Griesel G, Brink C, Baier PC, Capecchi MR, Mansouri A: In vivo and in vitro tissue-specific expression of green fluorescent protein using the cre-lox system in mouse embryonic stem cells. Stem Cells 2005, 23:10–15.

2. Collombat P, Xu X, Ravassard P, Sosa-Pineda B, Dussaud S, Billestrup N, Madsen OD, Serup P, Heimberg H, Mansouri A: The ectopic expression of Pax4 in the mouse pancreas converts progenitor cells into alpha and subsequently beta cells. Cell 2009, 138:449–462.

3. Dor Y, Brown J, Martinez OI, Melton DA: Adult pancreatic beta-cells are formed by self-duplication rather than stem-cell differentiation. Nature 2004, 429:41–46.

4. Gannon M, Ables ET, Crawford L, Lowe D, Offield MF, Magnuson MA, Wright CV: pdx-1 function is specifically required in embryonic beta cells to generate appropriate numbers of endocrine cell types and maintain glucose homeostasis. Dev Biol 2008, 314:406–417.

5. Lange A, Gegg M, Burtscher I, Bengel D, Kremmer E, Lickert H: Fltp(T2AiCre): a new knock-in mouse line for conditional gene targeting in distinct mono- and multiciliated tissues. Differentiation 2012, 83:S105–113.

6. Zhong F, Jiang Y: Endogenous Pancreatic beta Cell Regeneration: A Potential Strategy for the Recovery of beta Cell Deficiency in Diabetes. Front Endocrinol (Lausanne) 2019, 10:101.

7. Murtaugh LC: Pancreas and beta-cell development: from the actual to the possible. Development 2007, 134:427–438.

8. Veres A, Faust AL, Bushnell HL, Engquist EN, Kenty JH, Harb G, Poh YC, Sintov E, Gurtler M, Pagliuca FW, et al: Charting cellular identity during human in vitro beta-cell differentiation. Nature 2019, 569:368–373.

9. Ramond C, Beydag-Tasoz BS, Azad A, van de Bunt M, Petersen MBK, Beer NL, Glaser N, Berthault C, Gloyn AL, Hansson M, et al: Understanding human fetal pancreas development using subpopulation sorting, RNA sequencing and single-cell profiling. Development 2018, 145.

10. Augsornworawat P, Maxwell KG, Velazco-Cruz L, Millman JR: Single-Cell Transcriptome Profiling Reveals beta Cell Maturation in Stem Cell-Derived Islets after Transplantation. Cell Rep 2020, 32:108067.

11. Trapnell C, Cacchiarelli D, Grimsby J, Pokharel P, Li S, Morse M, Lennon NJ, Livak KJ, Mikkelsen TS, Rinn JL: The dynamics and regulators of cell fate decisions are revealed by pseudotemporal ordering of single cells. Nat Biotechnol 2014, 32:381–386.

12. Campbell KR, Yau C: Uncovering pseudotemporal trajectories with covariates from single cell and bulk expression data. Nat Commun 2018, 9:2442.

13. Wolf FA, Hamey FK, Plass M, Solana J, Dahlin JS, Gottgens B, Rajewsky N, Simon L, Theis FJ: PAGA: graph abstraction reconciles clustering with trajectory inference through a topology preserving map of single cells. Genome Biol 2019, 20:59.

14. Lareau CA, Ludwig LS, Muus C, Gohil SH, Zhao T, Chiang Z, Pelka K, Verboon JM, Luo W, Christian E, et al: Massively parallel single-cell mitochondrial DNA genotyping and chromatin profiling. Nat Biotechnol 2020.

15. Ludwig LS, Lareau CA, Ulirsch JC, Christian E, Muus C, Li LH, Pelka K, Ge W, Oren Y, Brack A, et al: Lineage Tracing in Humans Enabled by Mitochondrial Mutations and Single-Cell Genomics. Cell 2019, 176:1325–1339 e1322.

16. Enge M, Arda HE, Mignardi M, Beausang J, Bottino R, Kim SK, Quake SR: Single-Cell Analysis of Human Pancreas Reveals Transcriptional Signatures of Aging and Somatic Mutation Patterns. Cell 2017, 171:321–330 e314.

17. Courtney M, Gjernes E, Druelle N, Ravaud C, Vieira A, Ben-Othman N, Pfeifer A, Avolio F, Leuckx G, Lacas-Gervais S, et al: The inactivation of Arx in pancreatic alpha-cells triggers their neogenesis and conversion into functional beta-like cells. PLoS Genet 2013, 9:e1003934.

18. Talchai C, Xuan S, Lin HV, Sussel L, Accili D: Pancreatic beta cell dedifferentiation as a mechanism of diabetic beta cell failure. Cell 2012, 150:1223–1234.

19. van Arensbergen J, Garcia-Hurtado J, Moran I, Maestro MA, Xu X, Van de Casteele M, Skoudy AL, Palassini M, Heimberg H, Ferrer J: Derepression of Polycomb targets during pancreatic organogenesis allows insulin-producing beta-cells to adopt a neural gene activity program. Genome Res 2010, 20:722–732.

20. Marquina-Sanchez B, Fortelny N, Farlik M, Vieira A, Collombat P, Bock C, Kubicek S: Single-cell RNA-seq with spike-in cells enables accurate quantification of cell-specific drug effects in pancreatic islets. Genome Biol 2020, 21:106.

