## Supplemental information of the manuscript for "Single-cell transcriptome lineage tracing of human pancreatic development identifies distinct developmental trajectories of alpha and beta cells"

### Supplementary information for : A computational study of human pancreatic islet based on single cell RNA-seq data reveals distinct lineage features of alpha and beta cells

Li Lin [1, *]
Yufeng Zhang [2, *]
Weizhou Qian [1, *]
Yao Liu [3]
Yingkun Zhang [1]
Fanghe Lin [1]
Cenxi Liu [2]
Guangxing Lu [2]
YanLing Song [1]
Jia Song [4, #]
Chaoyong Yang [1, 4, #]
Jin Li [2, #]

[1] State Key Laboratory for Physical Chemistry of Solid Surfaces, Key Laboratory for Chemical Biology of Fujian Province, Key Laboratory of Analytical Chemistry, and Department of Chemical Biology, College of Chemistry and Chemical Engineering, Xiamen University, Xiamen, 361005, People's Republic of China

[2] State Key Laboratory of Genetic Engineering and School of Life Sciences, Fudan University, Shanghai, China.

[3] Department of Endocrinology and Metabolism, Shanghai Tenth People's Hospital, School of Medicine, Tongji University, Shanghai, China

[4] Institute of Molecular Medicine, Renji Hospital, School of Medicine, Shanghai Jiao Tong University, Shanghai 200127, China

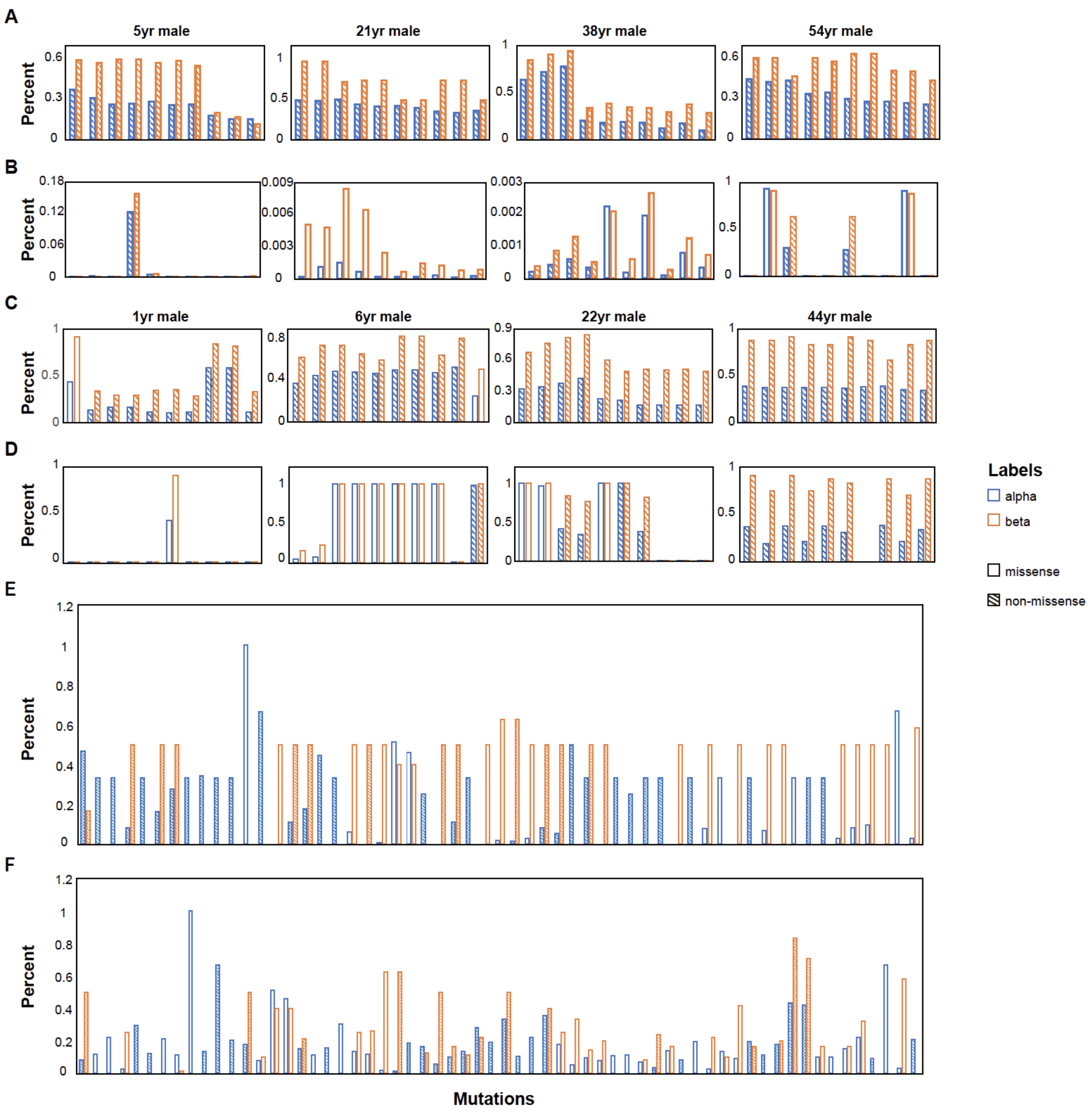


**Figure S1.** Nonsense and missense mutations in the top 10 high variance sites (A, C, E) or marker sites (B, D, F) of alpha and beta cells for primary cells (A-D) and iPS cell-derived cells (E-F).

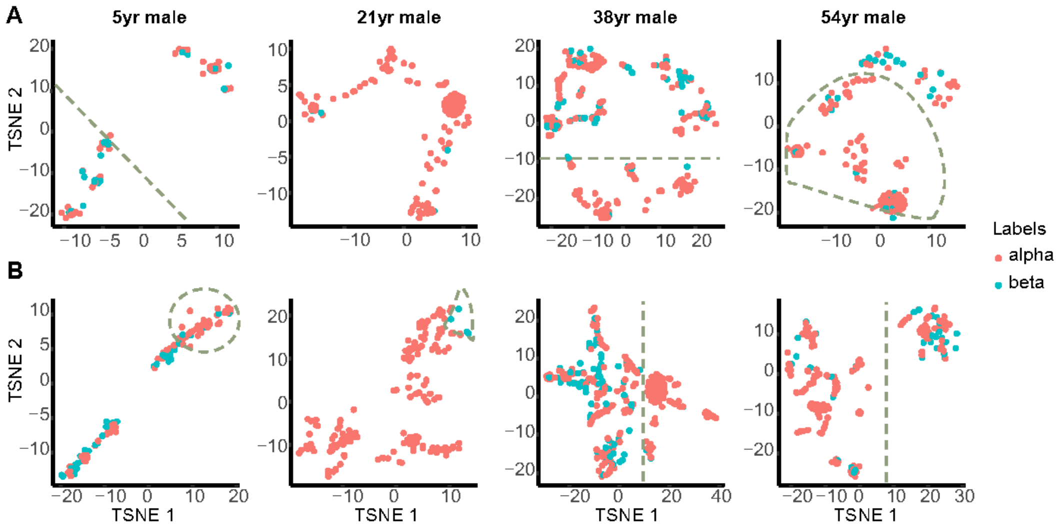


**Figure S2.** Clustering based on high variance sites (A) and marker sites (B) of human alpha and beta cells from four different donors.

**Table S1.** Summary of basic information of Smart-Seq2 dataset used in this study.

| ***Donors*** | ***1*** | ***2*** | ***3*** | ***4*** | ***5*** | ***6*** | ***7*** | ***8*** |
|---|---|---|---|---|---|---|---|---|
| *Age* | 21yr | 38yr | 1yr | 5yr | 44yr | 6yr | 54yr | 22yr |
| *Sex* | Male | Female | Male | Male | Female | Male | Male | Male |
| *Race* | Cau. | Cau. | African American | Cau. | African American | not provided | Cau. | Asian |
| *Cause of death* | Anoxia | Stroke | Anoxia | Auto accident | Stroke | Head trauma | Anoxia | Head trauma |
| *BMI* | 28.4 | 29.5 | 13.71 | 17.6 | 23.8 | (29kg) | 27.29 | 24.8 |
| *Number of alpha cells* | 157 | 226 | 38 | 61 | 141 | 71 | 171 | 133 |
| *Number of beta cells* | 4 | 69 | 48 | 35 | 24 | 43 | 36 | 89 |

1yr: "1 month old"

Cau.: Caucasian

**Table S2.** Summary of basic information of inDrops dataset used in this study.

| ***Protocol x1*** | | | |
|---|---|---|---|
| ***Stage*** | ***Stage4*** | ***Stage5*** | ***Stage6*** |
| *Number of alpha cells* | 2688 | 2402 | 3273 |
| *Number of nkx61 cells* | 1131 | ／ | ／ |
| *Number of neurog3 cells* | 451 | ／ | ／ |
| *Number of beta cells* | ／ | 226 | 233 |

**Table S3**. Properties of mutations used in the clustering of the Smart-Seq2 dataset.

| *Donor* | *High variance site* | | | *Marker site* | | |
|---|---|---|---|---|---|---|
| | *Site* | *Mutation* | *Mutation type* | *Site* | *Mutation* | *Mutation type* |
| 21yr_male | 16290 | C to T | Intergenic region | 8447 | A to T | Missense variant |
| | 16223 | C to T | Intergenic region | 15554 | C to T | Missense variant |
| | 16111 | C to T | Intergenic region | 15738 | A to C | Missense variant |
| | 16319 | G to A | Intergenic region | 5409 | A to T | Stop gained |
| | 16362 | T to C | Intergenic region | 11455 | C to A | Synonymous variant |
| | 12331 | A to C | Intragenic variant | 2383 | T to C | Intragenic variant |
| | 12335 | T to G | Intragenic variant | 6328 | C to T | Missense variant |
| | 16390 | G to A | Intergenic region | 6227 | T to C | Synonymous variant |
| | 16466 | A to C | Intergenic region | 7159 | T to C | Missense variant |
| | 12333 | A to T | Intragenic variant | 2056 | G to T | Intragenic variant |
| 38yr_female | 263 | A to G | Intergenic region | 2142 | A to G | Intragenic variant |
| | 16519 | T to C | Intergenic region | 2282 | C to T | Intragenic variant |
| | 16311 | T to C | Intergenic region | 825 | T to A | Intragenic variant |
| | 14743 | A to C | Intragenic variant | 2283 | C to T | Intragenic variant |
| | 14745 | C to G | Intragenic variant | 6690 | G to A | Stop gained |
| | 14744 | C to T | Intragenic variant | 6581 | A to G | Synonymous variant |
| | 14746 | A to G | Intragenic variant | 6134 | C to T | Synonymous variant |
| | 12336 | A to G | Intragenic variant | 2494 | C to T | Intragenic variant |
| | 14740 | A to T | Intragenic variant | 6698 | A to G | Synonymous variant |
| | 12331 | A to C | Intragenic variant | 9840 | T to C | Missense variant |
| 1yr_male | 14587 | A to G | Synonymous variant | 3484 | C to T | Missense variant |
| | 16519 | T to C | Intergenic region | 15734 | G to A | Missense variant |
| | 73 | A to G | Intergenic region | 4051 | G to A | Missense variant |
| | 146 | T to C | Intergenic region | 9277 | C to T | Missense variant |
| | 14744 | C to T | Intragenic variant | 10668 | G to A | Missense variant |
| | 14745 | C to G | Intragenic variant | 14587 | A to G | Synonymous variant |
| | 153 | A to G | Intergenic region | 2636 | G to A | Intragenic variant |
| | 8361 | G to T | Intragenic variant | 1998 | T to A | Intragenic variant |
| | 8362 | T to G | Intragenic variant | 2775 | A to G | Intragenic variant |
| | 14746 | A to G | Intragenic variant | 1836 | A to T | Intragenic variant |
| 5yr_male | 263 | A to G | Intergenic region | 1967 | T to G | Intragenic variant |
| | 16390 | G to A | Intergenic region | 1965 | A to G | Intragenic variant |
| | 16519 | T to C | Intergenic region | 15232 | A to T | Missense variant |
| | 14745 | C to G | Intragenic variant | 3121 | C to A | Intragenic variant |
| | 14746 | A to G | Intragenic variant | 2702 | G to A | Intragenic variant |
| | 14744 | C to T | Intragenic variant | 15270 | T to C | Missense variant |
| | 14740 | A to T | Intragenic variant | 14868 | T to A | Missense variant |
| | 12335 | T to G | Intragenic variant | 9901 | A to T | Missense variant |

| *Donor* | High variance site | | | Marker site | | |
|---|---|---|---|---|---|---|
| | *Site* | *Mutation* | *Mutation type* | *Site* | *Mutation* | *Mutation type* |
| 5yr_male | 12336 | A to G | Intragenic variant | 14831 | G to A | Missense variant |
| | 12333 | A to T | Intragenic variant | 9226 | C to G | Missense variant |
| 44yr_female | 16278 | C to T | Intergenic region | 16360 | C to T | Intergenic region |
| | 16286 | C to G | Intergenic region | 198 | C to T | Intergenic region |
| | 16311 | T to C | Intergenic region | 16311 | T to C | Intergenic region |
| | 16294 | C to T | Intergenic region | 189 | A to C | Intergenic region |
| | 16234 | C to T | Intergenic region | 16527 | C to T | Intergenic region |
| | 16360 | C to T | Intergenic region | 73 | A to G | Intergenic region |
| | 16527 | C to T | Intergenic region | 2033 | A to T | Intragenic variant |
| | 16145 | G to A | Intergenic region | 16278 | C to T | Intergenic region |
| | 16223 | C to T | Intergenic region | 186 | C to A | Intergenic region |
| | 16265 | A to C | Intergenic region | 16265 | A to C | Intergenic region |
| 6yr_male | 263 | A to G | Intergenic region | 13710 | A to T | Synonymous variant |
| | 73 | A to G | Intergenic region | 13710 | A to G | Synonymous variant |
| | 16362 | T to C | Intergenic region | 9007 | A to G | Missense variant |
| | 14745 | C to G | Intragenic variant | 10086 | A to G | Missense variant |
| | 14744 | C to T | Intragenic variant | 12705 | C to T | Synonymous variant |
| | 16223 | C to T | Intergenic region | 4769 | A to G | Synonymous variant |
| | 16124 | T to C | Intergenic region | 13105 | A to G | Missense variant |
| | 14740 | A to T | Intragenic variant | 5048 | T to C | Synonymous variant |
| | 16278 | C to T | Intergenic region | 2101 | C to T | Intragenic variant |
| | 10398 | A to G | Missense variant | 5773 | G to A | Intragenic variant |
| 54yr_male | 16362 | T to C | Intergenic region | 6222 | C to T | Missense variant |
| | 73 | A to G | Intergenic region | 4769 | A to G | Synonymous variant |
| | 15907 | A to G | Intragenic variant | 16092 | T to C | Intergenic region |
| | 16075 | T to C | Intergenic region | 15239 | T to A | Stop gained |
| | 16051 | A to G | Intergenic region | 15453 | T to A | Missense variant |
| | 16092 | T to C | Intergenic region | 16129 | G to C | Intergenic region |
| | 16129 | G to C | Intergenic region | 14983 | C to T | Synonymous variant |
| | 14744 | C to T | Intragenic variant | 2120 | G to A | Intragenic variant |
| | 14745 | C to G | Intragenic variant | 5390 | A to G | Synonymous variant |
| | 217 | T to C | Intergenic region | 9690 | G to A | Missense variant |
| 22yr_male | 73 | A to G | Intergenic region | 4769 | A to G | Synonymous variant |
| | 16217 | T to C | Intergenic region | 5465 | T to C | Synonymous variant |
| | 16519 | T to C | Intergenic region | 16261 | C to T | Intergenic region |
| | 16261 | C to T | Intergenic region | 16217 | T to C | Intergenic region |
| | 146 | T to C | Intergenic region | 15746 | A to G | Missense variant |
| | 16189 | T to C | Intergenic region | 1438 | A to G | Intragenic variant |
| | 14744 | C to T | Intragenic variant | 16519 | T to C | Intergenic region |

| ***Donor*** | **High variance site** | | | **Marker site** | | |
|---|---|---|---|---|---|---|
| | ***Site*** | ***Mutation*** | ***Mutation type*** | ***Site*** | ***Mutation*** | ***Mutation type*** |
| 22yr_male | 14745 | C to G | Intragenic variant | 1913 | G to A | Intragenic variant |
| | 14746 | A to G | Intragenic variant | 14874 | T to C | Missense variant |
| | 14740 | A to T | Intragenic variant | 15174 | C to T | Missense variant |

**Table S4**. Properties of mutations used in the clustering of the inDrops dataset.

| ***High variance site*** | | | ***Marker site*** | | |
|---|---|---|---|---|---|
| ***Site*** | ***Mutation*** | ***Mutation type*** | ***Site*** | ***Mutation*** | ***Mutation type*** |
| 646 | C to A | Intragenic variant | 2118 | T to A | Intragenic variant |
| 2114 | C to A | Intragenic variant | 3445 | C to A | Missense variant |
| 2115 | T to A | Intragenic variant | 3566 | C to A | Missense variant |
| 2118 | T to A | Intragenic variant | 4987 | C to A | Missense variant |
| 2944 | C to A | Intragenic variant | 5721 | T to A | Intragenic variant |
| 3242 | G to A | Intragenic variant | 5821 | G to A | Intragenic variant |
| 3303 | C to A | Intragenic variant | 6283 | C to A | Missense variant |
| 4384 | T to A | Intragenic variant | 6899 | G to A | Synonymous variant |
| 5567 | T to A | Intragenic variant | 7604 | G to A | Missense variant |
| 5638 | T to A | Intragenic variant | 8330 | T to A | Intragenic variant |
| 7486 | G to A | Intragenic variant | 8340 | G to A | Intragenic variant |
| 7604 | G to A | Missense variant | 10047 | C to A | Intragenic variant |
| 8340 | G to A | Intragenic variant | 10458 | C to A | Intragenic variant |
| 9269 | C to A | Synonymous variant | 11585 | T to A | Missense variant |
| 10457 | T to A | Intragenic variant | 15221 | G to A | Missense variant |
| 10458 | C to A | Intragenic variant | 15223 | C to A | Missense variant |
| 10466 | C to A | Intragenic variant | 2993 | T to C | Intragenic variant |
| 12215 | T to A | Intragenic variant | 3323 | T to C | Missense variant |
| 12563 | T to A | Missense variant | 5819 | T to C | Intragenic variant |
| 14722 | T to A | Intragenic variant | 10470 | A to C | Start lost<br>Splice region variant |
| 14924 | T to A | Missense variant | 10471 | T to C | Start lost<br>Splice region variant |
| 15221 | G to A | Missense variant | 12375 | T to C | Synonymous variant |
| 15223 | C to A | Missense variant | 13427 | A to C | Missense variant |
| 15949 | G to A | Intragenic variant | 14718 | A to C | Intragenic variant |
| 2492 | G to C | Intragenic variant | 14721 | G to C | Intragenic variant |
| 5865 | T to C | Intragenic variant | 1622 | A to G | Intragenic variant |
| 10006 | A to C | Intragenic variant | 1676 | A to G | Intragenic variant |
| 12339 | A to C | Initiator codon variant<br>Splice region variant | 1680 | A to G | Intragenic variant |
| 13427 | A to C | Missense variant | 1688 | A to G | Intragenic variant |
| 14718 | A to C | Intragenic variant | 1945 | A to G | Intragenic variant |
| 15228 | T to C | Missense variant | 2500 | A to G | Intragenic variant |
| 1616 | A to G | Intragenic variant | 2595 | A to G | Intragenic variant |
| 1676 | A to G | Intragenic variant | 2945 | A to G | Intragenic variant |
| 2035 | T to G | Intragenic variant | 2946 | A to G | Intragenic variant |
| 2595 | A to G | Intragenic variant | 2958 | A to G | Intragenic variant |

| *High variance site* | | | *Marker site* | | |
|---|---|---|---|---|---|
| ***Site*** | ***Mutation*** | ***Mutation type*** | ***Site*** | ***Mutation*** | ***Mutation type*** |
| 2600 | A to G | Intragenic variant | 3320 | A to G | Missense variant |
| 4394 | C to G | Intragenic variant | 3577 | A to G | Missense variant |
| 4411 | A to G | Intragenic variant | 3917 | A to G | Missense variant |
| 5547 | A to G | Intragenic variant | 4474 | A to G | Missense variant |
| 5651 | C to G | Intragenic variant | 7772 | A to G | Missense variant |
| 5907 | T to G | Missense variant | 7780 | A to G | Synonymous variant |
| 7543 | A to G | Intragenic variant | 8192 | A to G | Missense variant |
| 9210 | A to G | Missense variant | 8296 | A to G | Intragenic variant |
| 12349 | A to G | Missense variant | 9531 | A to G | Missense variant |
| 14456 | A to G | Missense variant | 10028 | A to G | Intragenic variant |
| 16015 | T to G | Intragenic variant | 10936 | C to G | Missense variant |
| 3309 | A to T | Initiator codon variant<br>Splice region variant | 13691 | A to G | Missense variant |
| 3571 | C to T | Missense variant | 14805 | A to G | Missense variant |
| 3904 | C to T | Missense variant | 15046 | A to G | Synonymous variant |
| 4406 | A to T | Intragenic variant | 1944 | C to T | Intragenic variant |
| 7479 | G to T | Intragenic variant | 2124 | A to T | Intragenic variant |
| 12960 | A to T | Synonymous variant | 2208 | A to T | Intragenic variant |
| 14500 | A to T | Missense variant | 2617 | A to T | Intragenic variant |
| 14501 | A to T | Missense variant | 3289 | A to T | Intragenic variant |
| 14753 | C to T | Missense variant | 3492 | A to T | Missense variant |
| 14766 | C to T | Missense variant | 4474 | A to T | Missense variant |
| 14926 | A to T | Synonymous variant | 4986 | A to T | Missense variant |
| | | | 8366 | A to T | Start lost<br>Splice region variant |
| | | | 12299 | A to T | Intragenic variant |
| | | | 14766 | C to T | Missense variant |
| | | | 14926 | A to T | Synonymous variant |
| | | | 15957 | C to T | Intragenic variant |
